# Identification of common sequence motifs shared exclusively among selectively packed exosomal pathogenic microRNAs during rickettsial infections

**DOI:** 10.1101/2023.01.06.522907

**Authors:** Jiani Bei, Yuan Qiu, Diane Cockrell, Qing Chang, Sorosh Husseinzadeh, Changcheng Zhou, Angelo Gaitas, Xiang Fang, Yang Jin, Kamil Khanipov, Tais B. Saito, Bin Gong

**Affiliations:** Department of Pathology, University of Texas Medical Branch, Galveston, Texas 77555, USA; Laboratory of Bacteriology, Division of Intramural Research, NIAID-NIH, Hamilton, Montana 59840, USA; Department of Neurology, Icahn School of Medicine at Mount Sinai, New York, New York 10029, USA; Department of Neurology, University of Texas Medical Branch, Galveston, Texas 77555, USA; Pulmonary and Critical Care Medicine Division, Department of Medicine, Boston University Medical Campus, Boston, Massachusetts 02118, USA; Department of Pharmacology, University of Texas Medical Branch, Galveston, Texas 77555, USA

**Keywords:** exosome, microRNA, sequence motif, exosome biogenesis, endothelial barrier dysfunction, fluidic AFM, rickettsial infection

## Abstract

We previously reported that microRNA (miR)23a and miR30b are selectively sorted into rickettsia-infected, endothelial cell-derived exosomes (*R*-ECExos). Yet, the mechanism remains unknown. The number of cases of spotted fever rickettsioses has been increasing in recent years, and infections with these bacteria cause life-threatening diseases by targeting brain and lung tissues. Therefore, the aim of the present study is to continue to dissect the molecular mechanism underlying *R*-ECExos-induced barrier dysfunction of normal recipient microvascular endothelial cells (MECs), depending on their exosomal RNA cargos. Rickettsiae are transmitted to human hosts by the bite of an infected tick into the skin. In the present study we demonstrate that treatment with *R*-ECExos, which were derived from spotted fever group *R parkeri* infected human dermal MECs, induced disruptions of the paracellular adherens junctional protein VE-cadherin and breached the paracellular barrier function in recipient pulmonary MECs (PMECs) in an exosomal RNA-dependent manner. Similarly, we did not detect different levels of miRs in parent dermal MECs following rickettsial infections. However, we demonstrated that the microvasculopathy-relevant miR23a-27a-24 cluster and miR30b are selectively enriched in *R*-ECExos. Bioinformatic analysis revealed that common sequence motifs are shared exclusively among the exosomal, selectively-enriched miR23a cluster and miR30b at different levels. Taken together, these data warrant further functional identification and characterization of a single, bipartition, or tripartition among ACA, UCA, and CAG motifs that guide recognition of microvasculopathy-relevant miR23a-27a-24 and miR30b, and subsequently results in their selective enrichments in *R*-ECExos.

## Introduction

The number of cases of spotted fever group rickettsioses (SFGR) has been increasing(*1*). Rickettsiae are transmitted to human hosts by the bite of an infected tick(*2-4*). *Rickettsia* (*R*.*) rickettsii, R. conorii*, and *R. australis* can cause life-threatening diseases. While *R. parkeri* is not life-threatening in humans(*5-8*), the organism is used as a model for pathogenic SFGR in a low biosafety level (BSL2) setting(*9*). Microvascular endothelial cells (MECs) are the primary mammalian host target cells of SFGR infections(*10, 11*). The most prominent pathophysiological effect during SFGR infections is increased microvascular permeability, followed by vasogenic cerebral edema and non-cardiogenic pulmonary edema with potentially fatal outcomes(*12, 13*). Cellular and molecular mechanisms underlying the dysfunction of the microvascular endothelial barrier in rickettsiosis remains largely unknown(*14-18*). Typically, rickettsial infection is controlled by appropriate antibiotic therapy if diagnosed early(*17, 19*). However, rickettsial infections can cause nonspecific signs and symptoms rendering early clinical diagnosis difficult(*10, 20*). Untreated or misdiagnosed infections are frequently associated with severe morbidity and mortality(*17, 20*). Comprehensive understanding of rickettsial pathogenesis is urgently needed for the development of novel therapeutics.

Cell-to-cell communication is critical for maintaining homeostasis and responding quickly to environmental stimuli(*21-44*). Besides direct intercellular contact, this communication is often mediated by soluble factors that can convey signals to a large repertoire of responding cells either locally or remotely. Extracellular vesicles (EVs) transfer functional mediators to neighboring and distant recipient cells(*31, 45*). EVs are broadly classified into two categories, exosomes (Exos) and microvesicles (MVs), distinguished by the cell membrane of origin(*36, 46-60*). Exosomes and MVs are also referred to as small (50-150 nm) and large (130-1000 nm) EVs, respectively(*49*). Exos contain many types of biomolecules, including proteins and nucleic acids, which contribute to disease pathogenesis, and are being actively investigated.

We reported a functional role played by endothelial Exos (ECExos) during rickettsial infection, and showed that MECs efficiently take up Exos *in vivo* and *in vitro*(*61*). We found that rickettsia-infected, MEC-derived exosomes (*R*-ECExos) induced disruption of the tight junctional (TJ) protein ZO-1 and barrier dysfunction of human normal recipient brain microvascular endothelial cells (BMECs), depending on their exosomal RNA cargos(*61*).

Many, if not all, the effects of Exos on the recipient cell are mediated by exosomal microRNA (miR) cargos(*62*). Mounting evidence demonstrates that miRs are selectively sorted into Exos, **independent** of the parent cellular levels(*50, 52-54, 63, 64*). Our deep sequencing and stem-loop RT-qPCR (slRT-PCR) assays revealed that *R. parkeri* infection triggers selectively enriched miRs, including microRNA (miR)23a and miR30b, in human umbilical vein endothelial cell (HUVEC)-derived *R*-ECExos(*61*). However, the levels of miR127, miR451, and miR92a were stable between mock ECExos and *R*-ECExos(*61*). Similar to our previous study using *R*. conorii(*65*), we did not detect different levels of these miRs in parent cell samples(*61*), suggesting that selective miRs are sorted into *R*-ECExos after infection with rickettsiae. Yet, the mechanism(s) remain unknown.

Some miRs are highly conserved, localized as clusters in the genome, transcribed together physically, and show similar expression profiles in cells(*66*). The miR23a-27a-24 cluster is intergenic with its own promoter on chromosome 19(*67-70*). miR23a and miR27a have been documented to modulate EC barrier function by directly targeting TJs(*67, 71*) and adherens junctions (AJs)(*68, 72*), respectively. miR24(*67, 73*), similar to miR30b(*74, 75*), is involved in governing EC function. We recently reported(*76*) that (1) ECExos deliver functional RNA oligos to recipient BMECs in the presence of RNase; (2) exosomal miR23a directly targets TJs; and (3) ECExo-delivered synthetic complementary antisense oligos (ASOs) of miR23a (23aASO) show therapeutic potential against microvasculopathy following infection with rickettsiae *in vitro*. Yet, we don’t know the impact on AJ, one of elemental components of the MEC barrier, following exposure to *R*-ECExos with regard to expression levels of other members of the miR23a cluster in *R*-ECExos.

In the present studies, we observed that exposure to dermal MEC (DMEC)-derived *R*-ECExos impairs AJ and paracellular barrier functions in BMECs and pulmonary MECs (PMECs), depending on exosomal RNA cargos. Individual miR analyses suggested that rickettsial infection triggered the selective enrichment of members of miR23a cluster in ECExos, potentially causing microvasculopathy in recipient MECs. Given that specific sequence motifs in miRs are recognized by specific RNA-binding proteins (RBPs) during selective packaging into Exos(*46, 50-54, 63*), we further identified common sequence motifs shared exclusively among the exosomal, selectively-enriched miR23a cluster and miR30b. These findings may help to dissect the molecular mechanism(s) of selective enrichment of microvasculopathy-relevant miRs in Exos during rickettsial infection by identifying and characterizing the functional miR sequence motif(s) for guiding RBP-conducted selective sorting.

## Results

### 1. *R*-ECExos disrupted AJs in recipient BMECs

Rickettsiae are transmitted to a human host by the bite of an infected tick into the skin(*2-4*). Size-exclusion chromatography (SEC) was utilized to isolate ECExos from human DMEC culture media that had been passed through two 0.2 μm filters(*61*). We routinely evaluated sizes and morphologies of isolated particles using atomic force microscopy (AFM)(*61*) and Tunable Resistive Pulse Sensing nanoparticle tracking analysis (NTA)(*77*), and verified the size distribution of isolated EVs in the range of 50 nm to 150 nm, which is the expected size of Exos (**Fig. 1A, B**). *R*-ECExos were purified 72 hrs post-infection from a multiplicity of infection (MOI) of 10 *R. parkeri*-infected DMECs. Quantitative real-time PCR validated that no rickettsial DNA copies were detected in the *R*-ECExo material (**Fig. 1C**). Using immunoblotting assays that employed rabbit hyperimmune sera against heat-killed SFGR(*78, 79*), no bacterial surface antigens was detected (**Fig. 1D**). As we reported(*61*), our purified ECExos and *R*-ECExos were free of bacteria or bacterial DNA, were intact, and did not aggregate.

**Figure 1.**
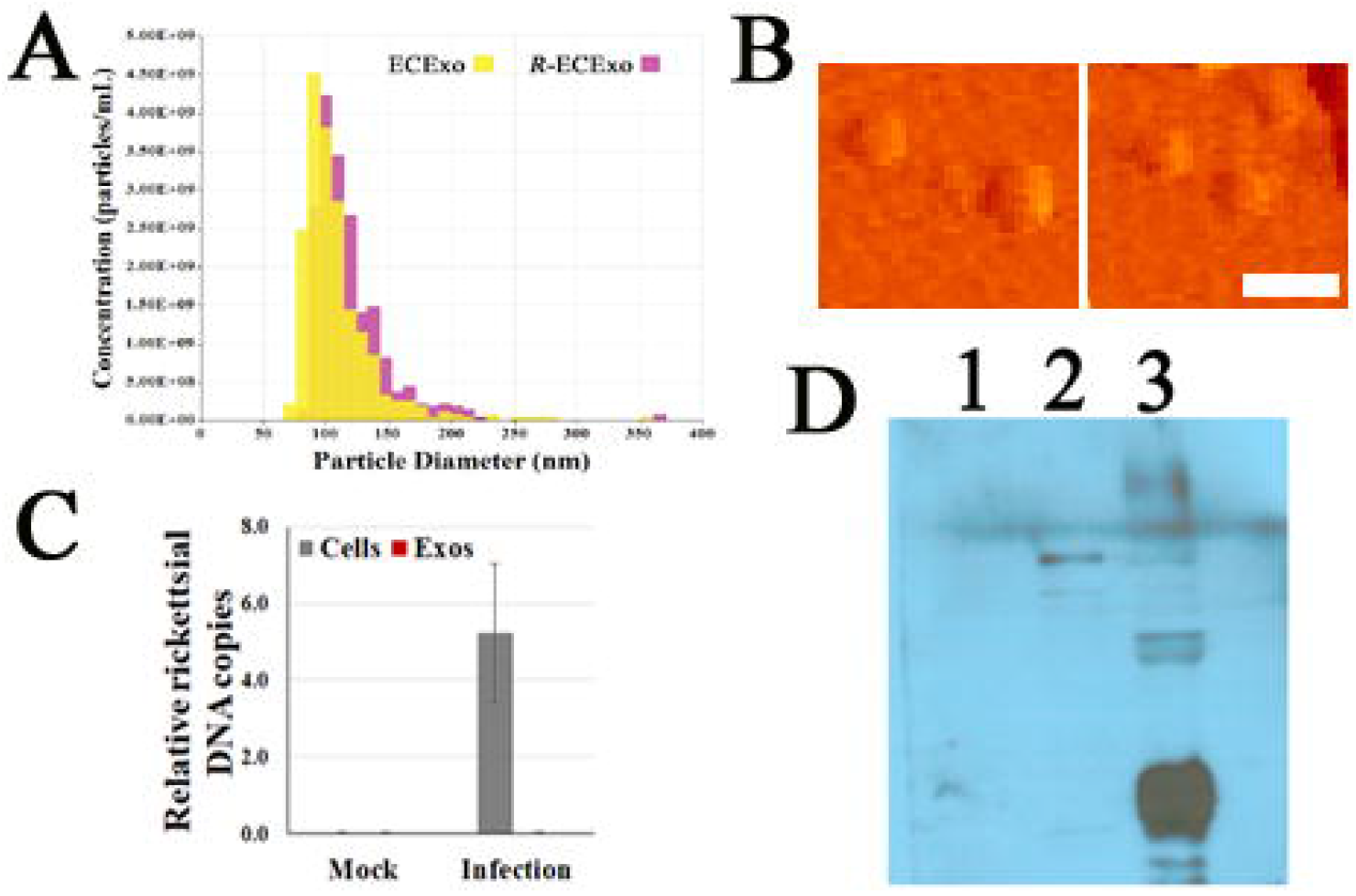
Characterization of DMEC-derived SEC-isolated ECExos. **(A)** Vesicle size distribution of isolated EVs was analyzed using NTA. **(B)** ECExo morphology was verified using AFM deflection image (scale bar, 200 nm). **(C)** Quantities of rickettsiae in parent DMECs and ECExos (*n*□=□3/group), determined by quantitative real-time PCR. Data are presented as means ± standard errors. **(D)** Bacterial surface antigens were examined in immunoblotting assay using 50 μg proteins extracted from 1-*R*-ECExos, 2-*R*-DMECs, and 3-rickettsial stocks using a rabbit hyperimmune sera against heat-killed SFG rickettsia(*78, 79*).

The endothelial barrier is tightest in the blood-brain barrier (BBB)(*80*). BBB dysfunctions underlie vasogenic cerebral edema during lethal SFGR infections(*13*), and can be caused by disruptions of TJs and adherens junctions (AJs), two elemental components of microvascular endothelial barrier(*80, 81*). We reported reduced or disorganized paracellular TJ protein ZO-1 in recipient BMECs of *R*-ECExos and demonstrated that *R*-ECExos induced dysfunction of normal recipient BMECs in an exosomal RNA cargo-dependent manner(*61, 76*). To evaluate the effect of *R*-ECExos on the AJs, we examined transmembrane adhesive protein VE-cadherin(*82*). We demonstrated that treatment with *R*-ECExos (1,000 particles/cell) induced disruptions of paracellular VE-cadherin, as evidence by reduced immunofluorescence staining of VE-cadherin (**Fig. 2A**).

**Figure 2.**
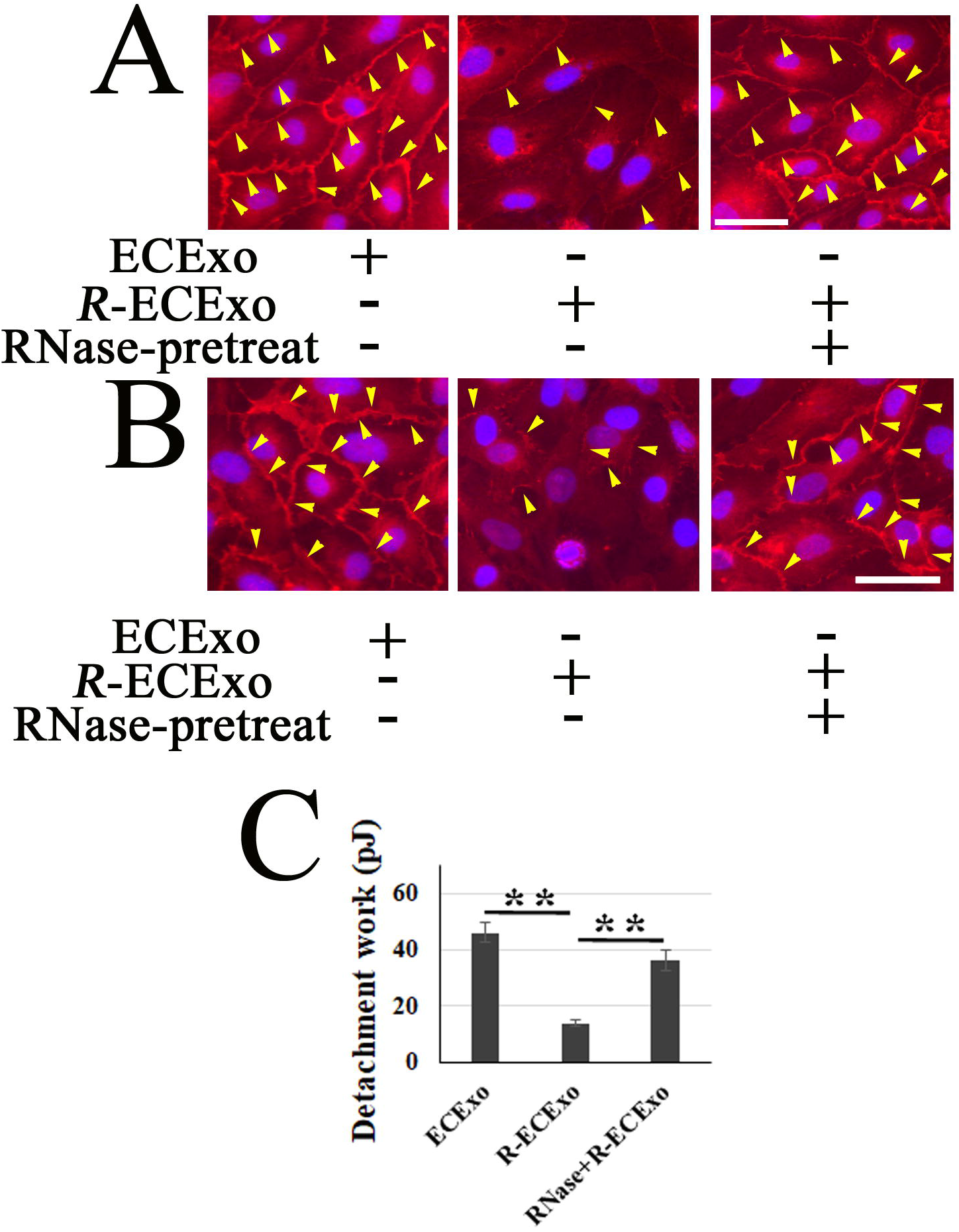
*R*-ECExos breach the paracellular barrier function in MECs, depending on their RNA cargos. **(A)** Representative immunofluorescence stainingof VE-cadherin in recipient BMECs after exposure to ECExos, *R*-ECExos, and *R*-ECExos that were treated with 20 μg/mL ribonuclease (RNase) in the presence or absence of 0.1% saponin as we reported(*61*). **(B)** Representative immunofluorescence stainingof VE-cadherin in recipient PMECs after exposure to ECExos, *R*-ECExos, and *R*-ECExos that were treated with 20 μg/mL ribonuclease (RNase) in the presence or absence of 0.1% saponin as we reported for ECExos from BMECs(*61*). **(C)** The lateral binding forces (LBFs) were assessed by Fluid AFM-measured detachment works in recipient PMECs after exposure to ECExos, *R*-ECExos, and *R*-ECExos that were treated with 20 μg/mL ribonuclease (RNase) in the presence or absence of 0.1% saponin as reported(*76, 85*).

### *2. R*-ECExos impair the paracellular barrier function in PMECs, depending on their RNA cargos

The PMEC barrier is relatively tight(*80*). EC-specific deletion of VE-cadherin gene *Cdh5* results in enhanced basal transvascular fluid flux in the lung, but not in brain tissue(*83, 84*). To evaluate the impact of VE-cadherin by exposure to *R*-ECExos, paracellular VE-cadherin in recipient PMECs was visualized, displaying suppression/disorganization as assessed by immunofluorescence staining (**Fig. 2B**). Furthermore, measuring the lateral binding forces (LBFs) between PMECs, the biomechanical nature of the endothelial barrier(*76, 85*), suggested that PMEC paracellular barrier dysfunction occurs after treatment with *R*-ECExos (**Fig. 2C**). However, the impact could be blocked if *R*-ECExos were pre-treated with RNase using the protocol as we reported(*61*).

These data suggest that *R*-ECExos impair normal recipient paracellular barrier function in PMECs in an exosomal RNA-dependent manner.

### 3. The miR23a-27a-24 cluster is selectively enriched in *R*-ECExos

We reported that miR30b and miR23a are selectively sorted into *R*-ECExos following *R. parkeri* infection(*61*). Using stem-loop RT-qPCR (slPCR), which is a common method for detecting small noncoding RNAs (sncRNAs) in EVs(*46, 55, 61*), we measured the expression levels of members of the miR23a cluster, miR27a and miR24, and EC-specific miR126(*86*) and EC-dominant miRlet7(*87*) in both ECExos and parent DMECs. We observed that, similar to miR23a, infections with *R. parkeri* trigger robust enrichment of miR27a and miR24 in *R*-ECExos while the exosomal levels of miR126 and let7 were stable (**Fig. 3**). Similarly(*65*), we did not detect different levels of these miRs in parent cells following rickettsial infections. These data suggest that members of the miR23a cluster and miR30b are selectively enriched in human *R*-ECExo.

**Figure 3.**
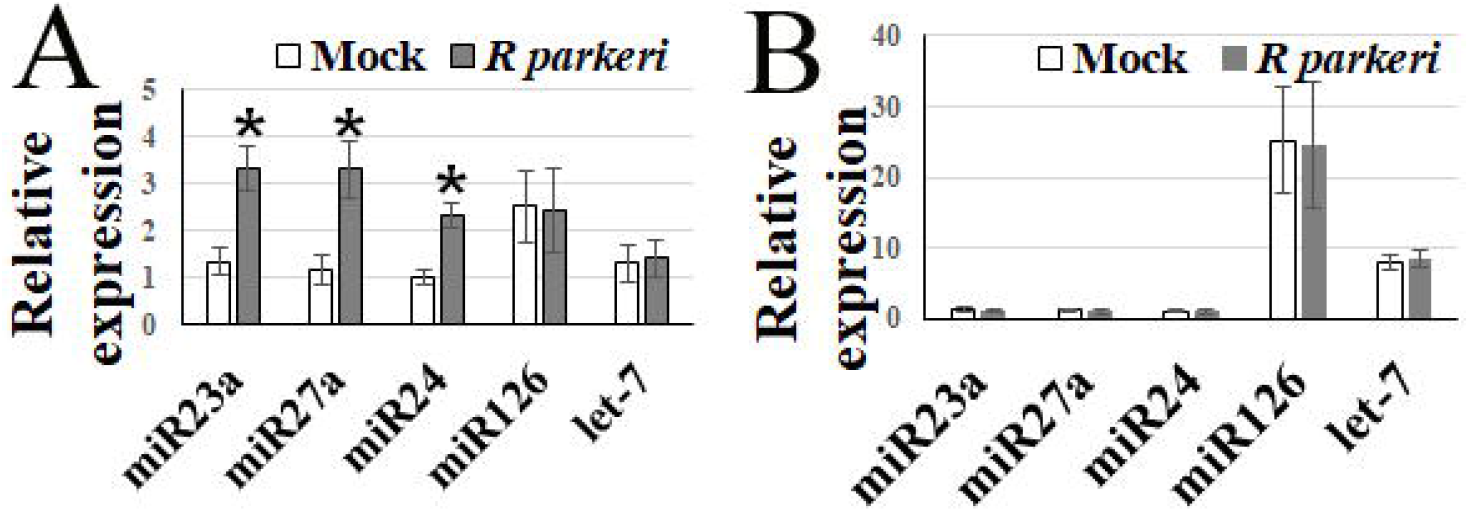
Members of the miR23a cluster are selectively enriched in *R*-ECExo. **(A)** Stem-loop RT-qPCR (slPCR) analysis of miRs obtained from ECExos (mock) and *R*-ECExos (*R* parkeri). * *P* < 0.05. **(B)** slPCR analysis of miRs obtained from normal (mock) and one-way analysis of variant rickettsial-infected parent DMECs. Statistical significance was determined using the one-way analysis of variance technique.

### 4. Common sequence motifs are shared exclusively among the exosomal, selectively-enriched miR23a cluster and miR30b

A subset of selective miRs enriched in Exos harbors a common sequence motif that is endowed with a functional capacity(*50, 52, 53, 63*). To identify the potential functional motifs shared among members of the miR23a cluster and miR30b, first we determined motif manipulation functions using R-4.2.2 (*cran*.*r-project*.*org*) and universal motif (*github*.*com/bjmt/universalmotif*) programs to create sequence logo of repeating patterns(*52, 53, 63*) among all slPCR-confirmed exosomal miRs, including 4 selective and 6 non-selective miRs (**Fig. 4A**). We further performed a motif analysis and found that there are different 3-to-5 nucleotide (nt)-long sequence motifs shared exclusively between members of miR23a cluster and miR30b (**Fig. 4B**). Notably, the 3 nt-long motifs ACA, UCA, and CAG are contained in miR23a-27a-24 and miR30b, but absent in 6 non-selective miRs in *R*-ECExos, suggesting these selective miRs harbor sequence motifs to guide unidentified exosomal RBP-mediated enrichment in Exos upon infection with tickettsiae.

**Figure 4.**
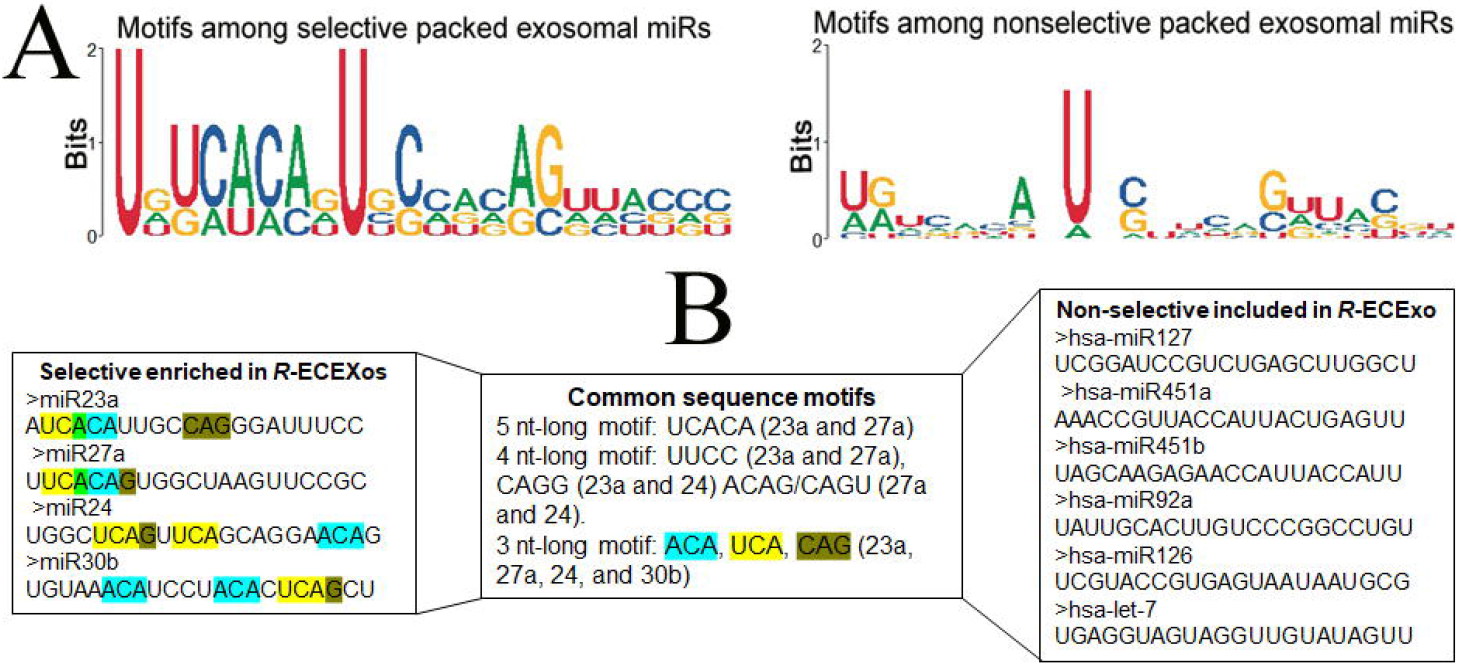
Bioinformatic identification of common sequence motifs shared exclusively among selectively enriched miRs in *R*-ECExos. **(A)** The motif manipulation functions were determined by sequence logo analysis of repetitive motifs among all slPCR-confirmed exosomal miRs, including 4 selective and 6 non-selective miRs. **(B)** Common sequence motifs shared among exosomal miRs at different levels were determined using motif analysis tools.

Identification of specific motifs in miR sequences recognized by specific RBPs during selective package into Exos(*46, 50-54, 63*), coupled with this preliminary information regarding selective enrichment of miR23a cluster and miR30b in ECExos during infection with rickettsiae, encourage us to characterize the molecular mechanism(s) for guiding the RBP-conducted miR23a cluster and miR30b selective sorting into *R*-ECExos in the future.

## Discussion

MECs are the primary targets of infection, and edema resulting from endothelial barrier dysfunction occurs in the brain and lungs in most cases of lethal SFGR infections in humans(*65, 88*). Capillaries provide the expansive surface area for efficient exchange of oxygen, carbon dioxide, and metabolites between blood and tissues(*80*). Nonfenestrated continuous endothelium is a major cellular component of capillaries in the brain and lung(*80, 89*). Endothelial barrier functions vary largely among organs. The endothelial barrier is tightest in the BBB(*80*), which restricts transendothelial flux of solutes and macromolecules and is attributed to junction tightness and limited transcytosis of specific molecules by transporter/receptor-mediated transcytosis(*90, 91*). The brain endothelial TJ complex is an elaborate structure composed of the transmembrane TJ proteins (mainly occludins and claudins) and intracellular scaffold proteins (mainly ZO family members)(*92, 93*), which functionally couples with VE-cadherin to the cytoskeleton(*94*). BBB breakdown can result in cerebral edema, elevated intracranial pressure, impaired cognitive or motor function, and can even be lethal(*93*). We reported reduced or disorganized paracellular TJ protein ZO-1 in recipient BMECs of *R*-ECExos and demonstrated that *R*-ECExos weakened the LBFs of normal recipient BMECs in an exosomal RNA cargo-dependent manner(*61, 76*). Now we further observed that exposure to *R*-ECExos disrupt paracellular VE-cadherins, although the underlying mechanism remains unknown.

The endothelial barrier is relative tight in lung tissue, and leakage can depress gas exchange and lead to hypoxia, hypercapnia, and even death(*80*). Disruption of VE-cadherin has been well documented underlying lung microvascular leakage and pulmonary edema in various contexts of diseases or pathologies(*95*). VE-cadherin connects adjacent ECs through homophilic interactions(*82*) and plays a functional role in an organ-specific manner(*80*). Conditional deletion of VE-cadherin gene *Cdh5* in ECs results in enhanced basal transvascular fluid flux in the lung and heart, but not brain and skin(*83, 84*). In the present study, we found that *R*-ECExo treatment induced suppression of paracellular VE-cadherin in recipient PMECs and impaired the paracellular barrier function through weakened LBFs, depending on their RNA cargos. Detailed mechanisms remain to be addressed.

Paralogous miRs, which have similar sequences but differ in their chromosomal localization, always show the same tendency to be localized in Exos. In contrast, mature miR complementary chains derived from the same pre-miRNA, which are therefore expressed at similar levels but have different sequences, can differ in their tendency for exosomal enrichment, suggesting that the sequences of mature miRs are important for determining their sorting into Exos(*46, 49-54, 63*). Some miRs are highly conserved, localized as clusters in the genome, transcribed together physically, and show similar expression profiles in cells(*66*). Alignment of the miR sequences revealed a significant number of miR paralogs among the clusters(*66*). There are two copies of the miR23-27-24 cluster in the genome, miR23a-27a-24 being intergenic with its own promoter on chromosome 19 and miR23b-27b-24 being intronic on chromosome 9(*67-70*). Now we show that miR23a/27a/24, similar to miR23a and miR30b in HUVEC-derived ECExos(*61*), were up-regulated in DMEC-derived *R*-ECExos 72 hrs post-infection (p.i.) with 10 MOI *R. parkeri*, while miR126 and miRlet-7 are stable (**Fig. 3**). Similar to our previous study(*61, 65*), we did not detect different levels of all these miRs in parent cells upon *R* infections, collectively suggesting the miR23a-27a-24 cluster and miR30b are selectively sorted into *R*-ECExos following *R*. infection. miR23a and miR27a have been documented to modulate EC barrier function by directly targeting TJs(*67, 71*) and AJs(*68, 72*), respectively; miR24(*67, 73*) and miR30b(*74, 75*) are involved in governing EC function, suggesting selectively enriched exosomal miRs cause microvasculopathy during rickettsial infections.

Alignment of the exosomal miR sequences suggested that the sequences of mature miRs are important for determining their sorting into Exos(*46, 49-54, 63*). A subset of selective miRs enriched in Exos harbors a common sequence motif that is endowed with a functional capacity(*50, 52, 53, 63*). These studies suggested that a single partition, bipartition, or tripartition among identified sncRNA sequence motifs directs recognition of specific sncRNAs resulting in their selective enrichment in Exos. Temoche-Diaz *et al*. reported that a bipartition (including motifs UUUG at the 3′ terminus and UGGA at the 5′ terminus), or possibly more complex motifs, directs miR122 selective sorting into breast cancer cell-derived Exos(*49*). A new remarkable report identified multiple special miR sequence motifs that control cell-type selective enrichment of miRs in Exos(*63*). Yet, information in the context of infections remain unknown. In the present study, we identified *in silico* different 3-to-5 nucleotide-long sequence motifs shared between the miR23a-27a-24 cluster and miR30b. Notably, the 3 nt-long motifs ACA, UCA, and CAG are contained in miR23a-27a-24 and miR30b, but absent in 6 non-selective miRs in R-ECExos. This prompted us to plan experiments using mutagenesis methods(*49, 50*) to characterize a single, bipartition, or tripartition among ACA, UCA, and CAG motifs-guided recognition of miR23a-27a-24 and miR30b, which subsequently results in their selective enrichment in *R*-ECExos following infection by SFGR.

Taken together, these data warrant further investigation of the mechanisms underlying selective packaging of specific exosomal miRs and suggests that cell-type exosomally targeting specific MVP miRs stabilizes EC function during infection with *R*, thus providing a novel strategy against rickettsioses.

## Materials and Methods

### Exo isolation and purification(*61*)

After *R. parkeri* at a MOI of 10 or mock infection for 72 hours, 11 mL of media from PMECs in T75 flasks was collected and filtered twice using 0.2 μm syringe filters. Ten mL of filtered media was placed in the qEV10 column (Izon, Medford, MA) for Exo isolation according to the manufacturer’s instructions. Number 7 through 10 fractions were collected and pooled as the Exo-enriched fractions, which were concentrated using 100,000 MWCO PES Vivaspin centrifugal filters (Thermo Fisher Scientific, Waltham, MA). All isolated bacteria-free Exos samples were routinely confirmed using plaque assay as we described(*96*). Conditioned culture media from PMECs that were infected with *R. parkeri* for 72 hours were used as positive control prior to the filtration. Nanoparticle tracking analysis (NTA) was performed using a qNano Gold system (Izon Science, Bellarie, TX) to determine the size and concentration of extracellular vesicle particles of each Exo sample. Exo samples (in 200 μl PBS) were stored at -80□ for further use.

### Nanoparticle tracking analysis

NTA was performed using the TRPS technique and analyzed on a qNano Gold system (Izon Science) to determine the size and concentration of EV particles. With the qNano instrument, an electric current between the two fluid chambers is disrupted when a particle passes through a nanopore NP150 with an analysis range of 70 nm to 420 nm, causing a blockade event to be recorded. The magnitude of the event is proportional to the amount of particles traversing the pore, and the blockade rate directly relates to particle concentration that is measured particle by particle. The results can be calibrated using a single-point calibration under the same measurement conditions used for EV particles (stretch, voltage, and pressure)(*97*). CPC200 calibration particles (Izon Science) were diluted in filtered Izon-supplied electrolyte at 1:500 to equilibrate the system prior to measuring EVs and for use as a reference. Isolated extracellular vesicle samples were diluted at 0, 1:10, 1:100, and 1:1,000. For the measurements, a 35 μl sample was added to the upper fluid chamber and two working pressures, 5 mbar and 10 mbar, were applied under a current of 120 nA. Particle rates between 200 and 1500 particles/min were obtained. The size, density, and distribution of the particles were analyzed using qNano software (Izon Science).

### Imaging of label-free extracellular vesicles using atomic force microscopy (AFM)

Tapping mode AFM provides a three-dimensional image of surface structures, including height image or deflection image (*98-100*), and is commonly used to evaluate the integrity of EVs at the single particle level. To obtain deflection images, the purified and concentrated EV sample was diluted at 1:10, 1:100, and 1:1000 with molecular grade water. Glass coverslips were cleaned three times with ethanol and acetone, then three times with molecular grade water. The coverslip was coated with the diluted EV samples on the designated area for 30 minutes, before being examined using an AFM (CoreAFM, Nanosurf AG, Liestal, Switzerland) with the contact mode in the air. A PPP-FMR-50 probe (0.5-9.5N/m, 225μm in length and 28μm in width, NANOSENSORS, Neuchatel, Switzerland) was used. The parameters of the cantilever were calibrated using the default script from the CoreAFM program using the Sader *et al*. method(*101*). The cantilever was approached to the sample under the setpoint of 20 nN, and topography scanning was done using the following parameters: 256 points per line, 1.5 seconds per line in a 5-μm x 5-μm image.

### *In vitro* cell model of Exo treatment

Human BMECs or PMECs (iXCells, SanDiego, CA) were cultivated in 5% CO_2_ at 37°C on type I rat-tail collagen-coated round glass coverslips (12 mm diameter, Ted Pella, Redding, CA) or inserts in 24-well plates (0.4 μm polyester membrane, CoStar, Thermo Fisher Scientific, Rockford, IL) until 90% confluence was achieved(*102*). Cells were exposed to *R*-ECExos (1,000 particles/cell) for 6 hrs prior to treatment with exosomes or naked oligonucleotides (IDT) by direct addition of the specified quantity of exosomes or oligonucleotides in the culture media for 66 hrs. Cells were subjected to downstream assays, and all experiments were performed in triplicate.

### Fluidic AFM assay for measuring lateral binding forces (LBFs) of PMECs

The LBF is measured using the FluidFM system (Cytosurge, Nanosurf AG, Liestal, Switzerland). A micropipette fluidic AFM cantilever with an 8 μm aperture and spring constant of 2 N/m (CytoSurge) was calibrated using the Sader method in air and liquid by performing deflection and crosstalk compensation. Using an inverted microscope, the cantilever was kept at 20 mbar positive pressure by the fluidic pressure controller (Cytosurge) of the FluidFM system and manipulated to approach the surface of a PMEC in a monolayer. Prior to every measurement, the cantilever was set 45□μm away from the cell. The cantilever approached the target cell at 5□μm/sec until a set point of 20□nN was reached. After a set pause of 10 sec, suction pressure of - 200□mbar decreased to -400 mbar to ensure a seal between the apical surface of the cell and the aperture of the hollow cantilever. The cantilever was vertically retracted to a distance of 100 μm to separate the targeted PMEC from the substrate and the monolayer, and the unbinding effort was assessed by measuring the work done (in picojoules [pJ]), which was calculated by integrating the area under the force-distance (F-D) curve using software as we described(*96, 103*). The captured PMEC was released by applying 1,000 mbar positive pressure prior to moving the cantilever to another individual PMEC that did not contact other cells in the same culture. The LBF value was obtained using the method of Sancho *et al*. (*104*) by measuring the unbinding force/work required to separate the individual PMEC from the substrate. The Temperature Controller (NanoSurf) kept the liquid environment at 37°C during all measurements.

### Antibodies and other reagents

Anti-VE-cadherin rabbit antibodies were purchased from Abclonal (Woburn, MA). AlexaFluor 594-conjugated goat anti-rabbit IgG, and DAPI were purchased from Invitrogen (Carlsbad, CA). Normal rabbit IgGs were purchased from Agilent (Santa Clara, CA). Endothelial Cell Growth Medium and fetal bovine sera were obtained from Cell Applications, Inc. (San Diego, CA). Unless otherwise indicated, all reagents were purchased from Thermo Fisher Scientific.

### Western immunoblotting

For western immunoblotting, equal amounts of soluble protein were subjected to 10% SDS–polyacrylamide gel electrophoresis (SDS-PAGE). Proteins were transferred onto a polyvinylidene difluoride membrane and then incubated with primary antibody (1:5,000 for anti-SFGR antibodies) at 4□ overnight, followed by incubation with a secondary antibody at 1:10,000 for 2 hrs. A goat anti-rabbit IgG and IgM (H+L)-HRP (Thermo Fisher Scientific) was used as the secondary antibodies. Blots were visualized using the Pierce™ ECL2 Western Blotting Substrate kit (Thermo Fisher Scientific).

### Immunofluorescence (IF)

For IF of VE-cadherin in BMECs or PMECs, cells were incubated with anti-VE-cadherin rabbit polyclonal antibody (1:100) for 2 hrs, followed by AlexaFluor 594-conjugated goat anti-rabbit IgG (1:1000) for 30 min. Nuclei were counter-stained with DAPI. A rabbit polyclonal IgG (Thermo Fisher) served as a negative control (*107*). Fluorescent images were analyzed using an Olympus BX51 epifluorescence or Nikon A1R MP ECLIPSE T*i* confocal microscope with *NIS*-Elements imaging software (version 4.50.00) using a final 40x optical zoom.

### Statistical analysis and motif analysis

All data were presented as mean± standard error of the mean and analyzed using SPSS version 22.0 (IBM^®^ SPSS Statistics, Armonk, NY). A two-tailed Student’s *t*-test or one-way analysis of variance (ANOVA) was used to explore statistical differences. If the ANOVA revealed a significant difference, a post hoc Tukey’s test was further adopted to assess the pairwise comparison between groups. The level of statistical significance for all analyses was set at *p*<0.05.

The process of motif analysis demands a program to compare and evaluate various motifs, including their sequences and frequencies in a sequence sample. We wrote a program that can compare the motifs, including the functions of matching the orders of the motifs and counting the numbers of the occurrence of the motifs in a sample sequence. This program is coded to match the 3, 4, or 5 nucleotide-long sequence motif that were derived from sequence logo analysis of repeating patterns. First, to determine the templates, we defined the motifs into strings. Next, the sample sequence is thoroughly scrutinized for the various sizes of motifs using these templates, with each match resulting in an increase in the count and the recorded positioning of the motif. Whenever a match is found, the program will add a count and record the relative positions. Finally, the identified motifs are validated through mapping back to the sample sequences.

## Acknowledgements

We thank Dr. Ulrike Munderloh, Dr. Rong Fang, and Dr. William Russell for discussions during the planning phases of the experiments. We gratefully acknowledge Dr. Kimberly Schuenke for her critical review and editing of the manuscript. This work was supported by NIH grant R01AI121012 (BG), R21AI137785 (BG), R21AI154211(BG), R03AI142406 (BG), R21AI144328 (BG), R21AG066060 (XF). The sponsors had no role in the study design, data collection and analysis, decision to publish, or preparation of the manuscript. This work was supported in part by the Division of Intramural Research, NIAID, NIH (TBS) and the Sealy and Smith Foundation (XF).

## Authorship Contributions

JB, YQ, DC, QC, SH, and CZ performed experiments. JB, YQ, QC, AG, KK analyzed data. YQ wrote the program for motif analysis. AG, XF, YJ, WR, and KK designed the study. TS and BG designed the study, performed experiments, analyzed data, and wrote the manuscript.

## Notes

### Competing Interest Statement

The authors have declared no competing interest.

